# A cytokine-responsive promoter is required for distal enhancer function mediating the hundreds-fold increase in milk protein gene expression during lactation

**DOI:** 10.1101/2023.02.06.527375

**Authors:** Hye Kyung Lee, Chengyu Liu, Lothar Hennighausen

**Affiliations:** Laboratory of Genetics and Physiology, National Institute of Diabetes and Digestive and Kidney Diseases, US National Institutes of Health, Bethesda, Maryland 20892, USA; Transgenic Core, National Heart, Lung, and Blood Institute, US National Institutes of Health, Bethesda, Maryland 20892, USA

**Author notes:** Correspondence to: H.K.L and L.H.

## Abstract

During lactation, specialized cells in the mammary gland produce milk to nourish the young. Milk protein genes are controlled by distal enhancers activating expression several hundred-fold during lactation. However, the role of promoter elements is not understood. We addressed this issue using the *Csn2* gene, which accounts for 10% of mRNA in mammary tissue. We identified STAT5 and other mammary transcription factors binding to three distal candidate enhancers and a cytokine-response promoter element. While deletion of the enhancers or the introduction of an inactivating mutation in a single promoter element had a marginable effect, their combined loss led to a 99.99% reduction of *Csn2* expression. Our findings reveal the essential role of a promoter element in the exceptional activation of a milk protein gene and highlight the importance of analyzing regulatory elements in their native genomic context to fully understand the multifaceted functions of enhancer clusters and promoters.

## Introduction

While enhancers control gene activation elicited by intra- and extra-cellular signals^1^, promoter elements are thought to control basal gene activity. Based on our previous studies, distal enhancers and super-enhancers (SE) control the exceptional expression of mammary genes during pregnancy and lactation^2–4^ (Lee et al., accompanying manuscript). However, it is not clear if this mechanism is operative across all pregnancy-regulated mammary genes and the importance of promoter-proximal elements in the cell-specific gene activation remains elusive. The five casein genes (*Csn1s1, Csn2, Csn1s2a, Csn1s2b* and *Csn3*) account for approximately 80% of milk proteins and 50% of total mRNA in mammary tissue at day 10 of lactation (L10)^5^, and their several hundred-fold induction during pregnancy is controlled by enhancers through the JAK-STAT signaling pathway^6–8^.

## Results

To gain insight into the possible contribution of promoter elements we dug deeper and investigated the regulation of the *Csn2* and *Csn1s2a* genes that are transcribed in opposite directions and whose promoters are separated by 75 kbp (Fig. 1). Both genes are activated up to 10,000-fold between non-parous and lactating mammary tissue and their combined mRNAs account for approximately 20% of total mRNA during lactation (Fig. 1a; Supplementary Table 1). The mammary transcription factors (TF) STAT5, glucocorticoid receptor (GR) and NFIB bound at five candidate enhancers (E1 to E5) that also coincided with activating histone marks (Fig. 1b). While all five enhancers are bound by STAT5, canonical recognition sites (GAS motifs) are present in only four, suggesting that binding to E3 is through another TF, possibly GR or NFIB. Of note, STAT5 binds to two distinct regions in E1, E2, and E4 suggesting additional internal complexity. STAT5 and GR binding was also associated with the *Csn2* promoter, coinciding with a canonical (TTCn3GAA) and a non-canonical (TTCn4GAA) GAS motif, as well as *Csn1s2a* (Fig. 1c). No bona fide GR binding site was detected suggesting GR binding through STAT5. The H3K27ac patterns differed between enhancer and promoter elements. While TF binding at enhancers was within a H3K27ac ‘gap’, binding at promoters was outside a classical gap (Fig. 1c). This points to the possibility that TF binding at enhancers and promoters serves a different purpose.

**Fig. 1.**
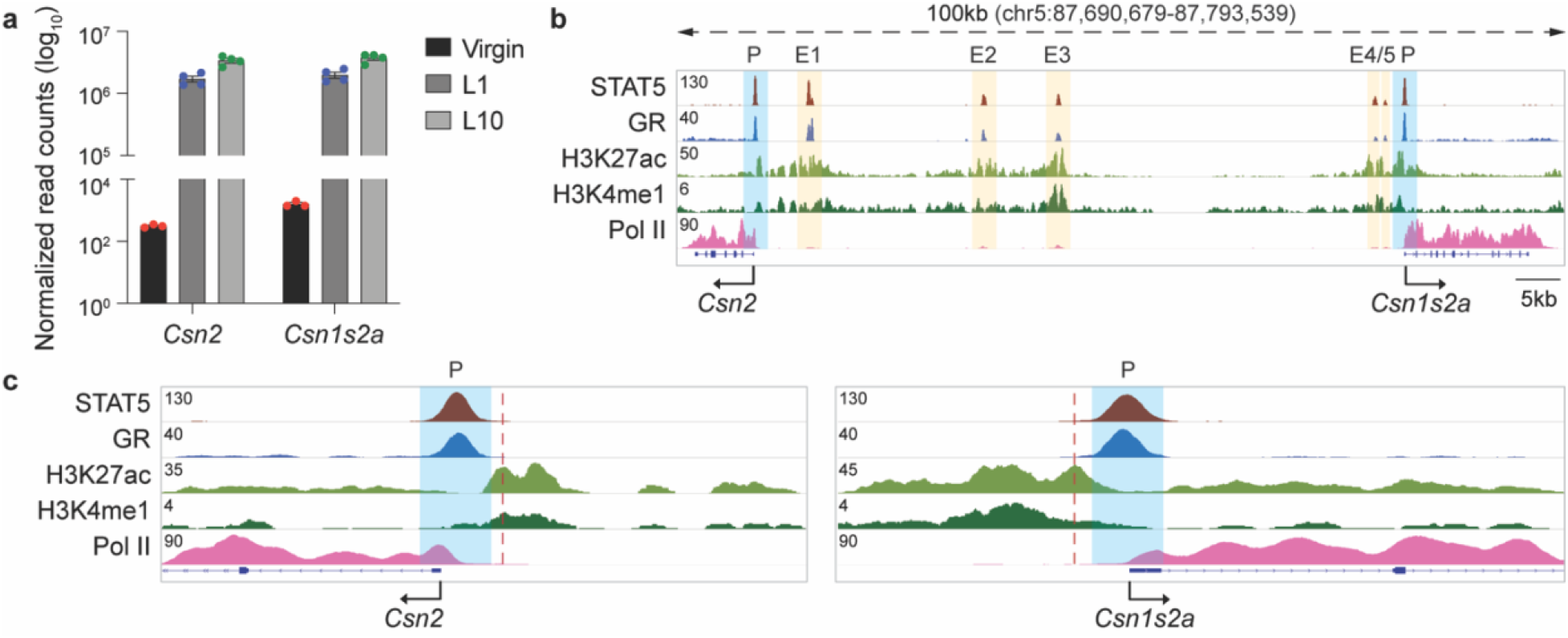
Characteristics of the *Csn2* locus. **a,** mRNA levels of *Csn2* and *Csn1s2a* genes were measured by RNA-seq at virgin, lactation day one (L1) and L10. (Virgin, *n* = 3; L1, L10, *n* = 4) **b,** Genomics features of the *Csn2* locus were identified by ChIP-seq data in lactating mammary tissue. **c,** ChIP-seq data at transcriptional start sites (TSS) provided structural information of regulatory elements of the *Csn2* and *Csn1s2a* genes at L1.

Next, we investigated the physiological role of the three candidate enhancers located upstream of *Csn2,* E1 (−6 kbp), E2 (−25 kbp) and E3 (−35 kbp) (Fig. 2a). All three candidate enhancers are bound by STAT5 and to a lesser extent by GR. We used CRISPR/Cas9 genome editing to delete these enhancers and monitor their functional significance in mammary tissue at day 1 of lactation (L1). Neither the deletion of E1 nor the combined deletion of E2 and E3 resulted in a significant reduction of *Csn2* mRNA levels (Fig. 2b). The combined deletion of all three enhancers resulted in a ~50% reduction of *Csn2* mRNA levels. The deletion of enhancers was validated using ChIP-seq analyses for activating histone marks, STAT5 and GR (Fig. 2c). Expression of *Csn1s2a* in L1 mammary tissue was not impaired significantly in these mutants.

**Fig. 2.**
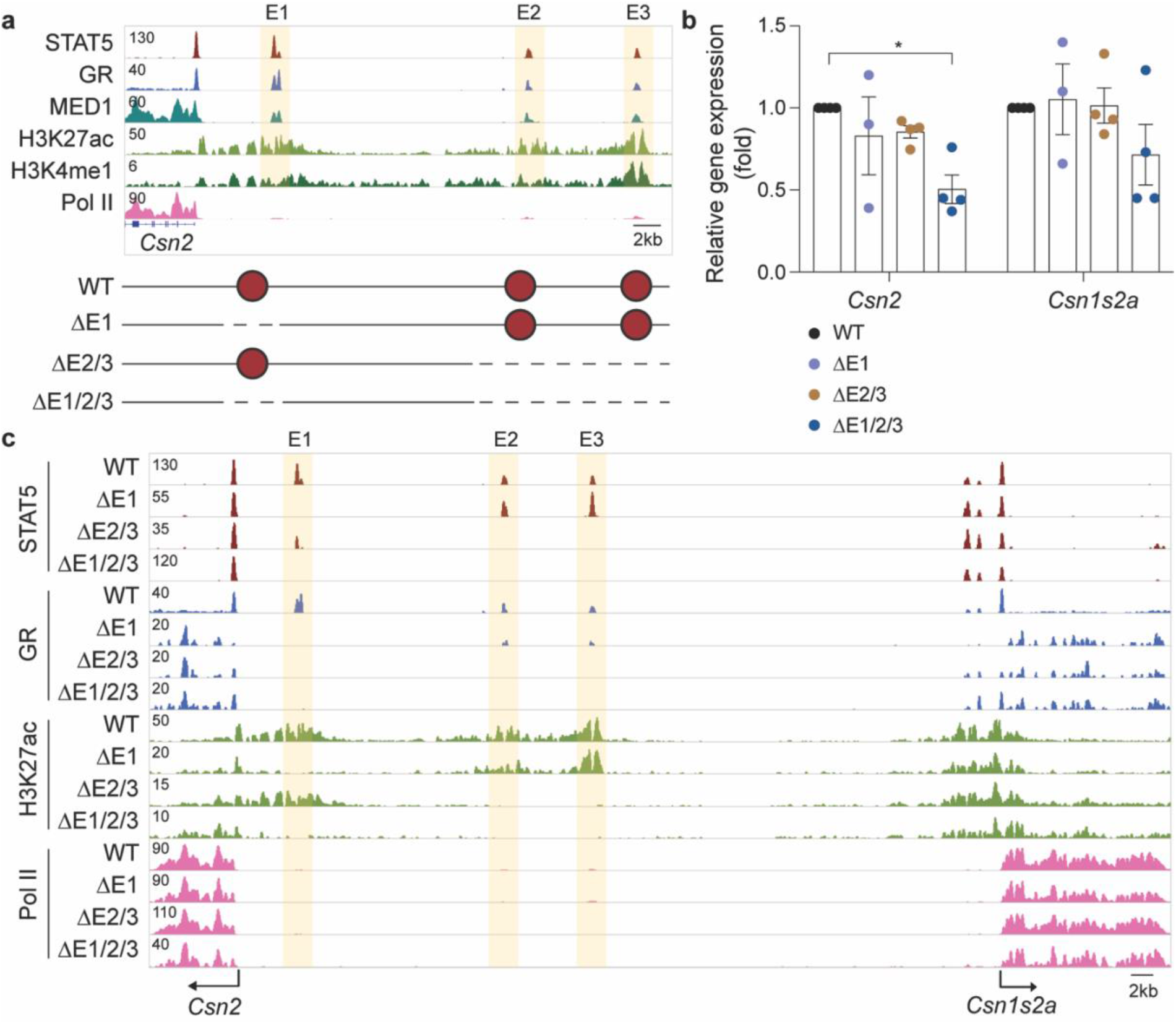
Activity of putative *Csn2* enhancers. **a,** The putative *Csn2* enhancers were identified by ChIP-seq for mammary transcription factors (TF) and activating histone marks at day one of lactation (L1). The presence of H3K27ac and H3K4me1 marks indicated three candidate enhancers, E1 at −6 kb, E2 at −25 kb and E3 at −35 kb. Diagram shows the enhancer deletions introduced in mice using CRISPR/Cas9 genome editing. Red circles mark the enhancers. **b,** Expression of the *Csn2* gene was measured in lactating mammary tissue (L1) from WT and mutant mice carrying enhancer deletions by qRT–PCR and normalized to *Gapdh* levels. Results are shown as the means ± s.e.m. of independent biological replicates (WT, ΔE2/3, ΔE1/2/3, *n* = 4; ΔE1, *n* = 3). One-way ANOVA followed by Dunnett’s multiple comparisons test was used to evaluate the statistical significance of differences between WT and each mutant mouse line. **c,** ChIP-seq analysis shows the genomic structure of *Csn2* locus in lactating mammary tissue of WT and mutant mice. The orange shades indicate enhancers.

Based on these results the three enhancers contribute only modestly to the activation of the *Csn2* gene, pointing to the presence of additional, possibly more prominent, regulatory elements. We therefore focused on the two STAT5 binding sites within the promoter region (Fig. 3a). We used deaminase base editing and introduced disabling point mutations into the palindromic sequences of the canonical and non-canonical GAS motifs located within 150 bp of the TSS. Mutations in both GAS sites (ΔP) resulted in an approximately 80% reduction of *Csn2* expression in L1 mammary tissue (Fig. 3b; Supplementary Table 2). These findings begged the question of potential synergy between the cytokine-responsive promoter elements and the distal enhancers. We addressed this in mice lacking the three enhancers and carrying point mutations in the promoter elements. The combined mutations resulted in an almost complete silencing of the *Csn2* gene, with expression levels reduced by three orders of magnitude (99.99%) (Fig. 3b). Loss of these elements coincided with the complete absence of Pol II coverage and H3K27ac (Fig. 3c). The three enhancers and the promoter-based STAT5 site are specifically active during pregnancy and their deletion did not affect expression in mammary tissue of virgin mice (Supplementary Fig. 1; Supplementary Table 3).

**Fig. 3.**
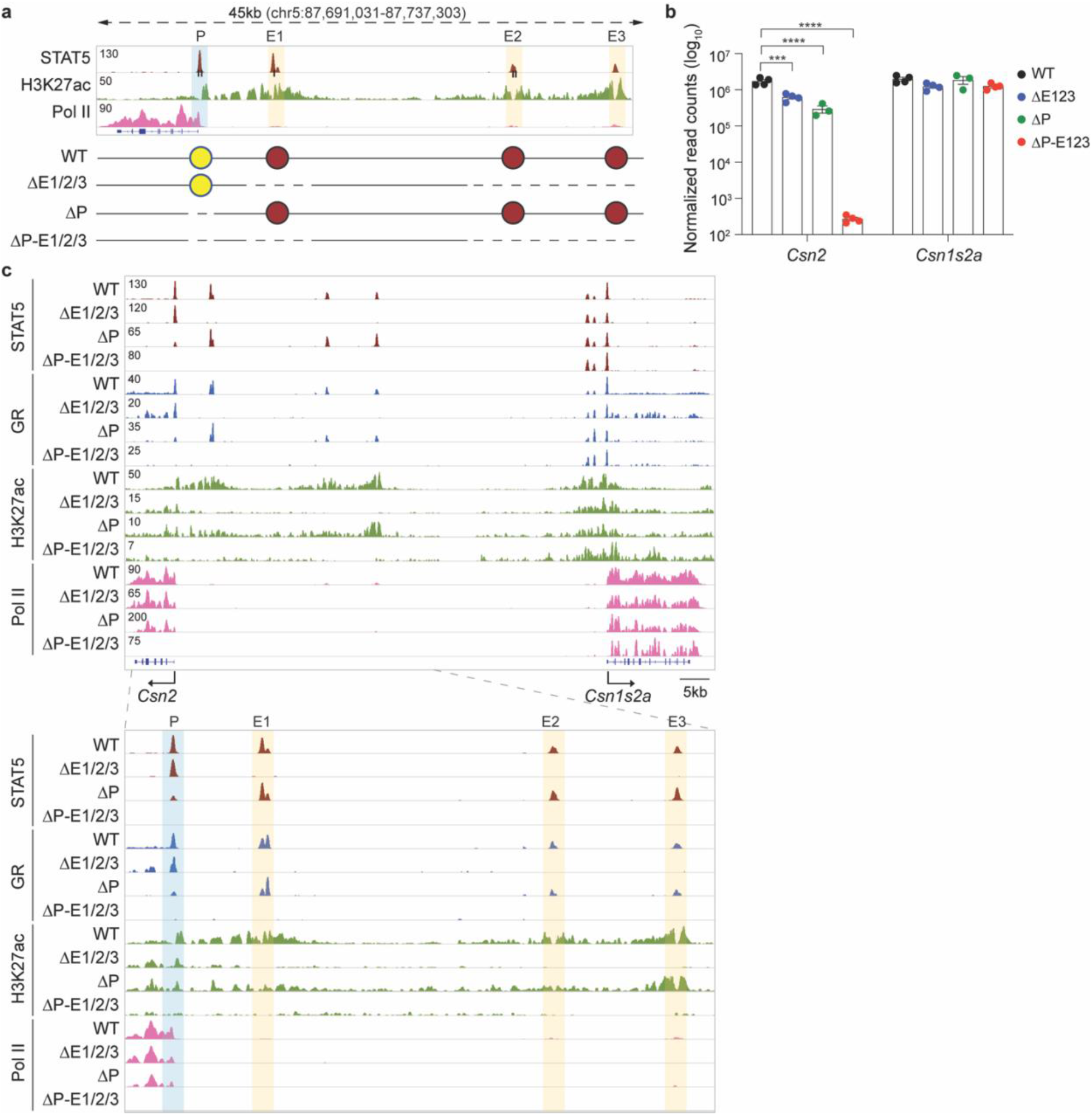
Synergy between promoter-based cytokine response elements and distal enhancers. **a,** The *Csn2* promoter region was characterized through ChIP-seq for STAT5, activating histone marks and Pol II loading at day one of lactation. Diagram shows the enhancer deletions and promoter mutations introduced in the mouse genome using CRISPR/Cas9 genome editing and deaminase base editing, respectively. **b,** *Csn2* mRNA levels were measured by RNA-seq in lactating mammary tissue isolated from WT mice and mice carrying disabling mutations in the two GAS motifs in *Csn2* promoter (ΔP) in the presence and absence of the three distal enhancers (ΔE1/2/3) (WT, ΔE1/2/3, ΔP-E1/2/3, *n* = 4; ΔP, *n* = 3). One-way ANOVA followed by Dunnett’s multiple comparisons test was used to evaluate the statistical significance of differences between WT and each mutant mouse line. **c,** Genomic features of the *Csn2* locus were investigated by ChIP-seq in lactating mammary tissue of WT, ΔE1/2/3, ΔP and ΔP-E1/2/3 mice. The highlighted blue and orange shades indicate a promoter and enhancers, respectively.

There is consensus that all seven STATs, except for STAT6, activate gene expression through canonical GAS motifs. A single STAT5 peak covers the closely spaced canonical and non-canonical GAS motifs in the *Csn2* promoter (Fig. 4) making it difficult to discern if either site or both are bound by STAT5 and convey transcriptional activity. To distinguish the relative significance of the two GAS motifs we introduced individual mutations, both in the presence and absence of the three distal enhancers (Fig. 4a). In the presence of the distal enhancers, mutation of the canonical site (A) did not impact *Csn2* expression during lactation, and a 50% reduction was observed upon the simultaneous deletion of the enhancers (ΔP-E1/2/3-A) (Fig. 4b). Introduction of a disabling mutation into the non-canonical site (B) resulted in a ~50% reduction of *Csn2* mRNA and the additional absence of the distal enhancers resulted in a complete loss of *Csn2* induction (ΔP-E1/2/3-B) and levels were four orders lower than in mice carrying intact regulatory elements. STAT5 binding was still observed upon loss of the canonical GAS motif and completely absent in the non-canonical mutant (Fig. 4c). These results provide evidence that the non-canonical GAS motif is a key element in the Csn2 promoter and STAT5 preferentially binds at the non-canonical site.

**Fig. 4.**
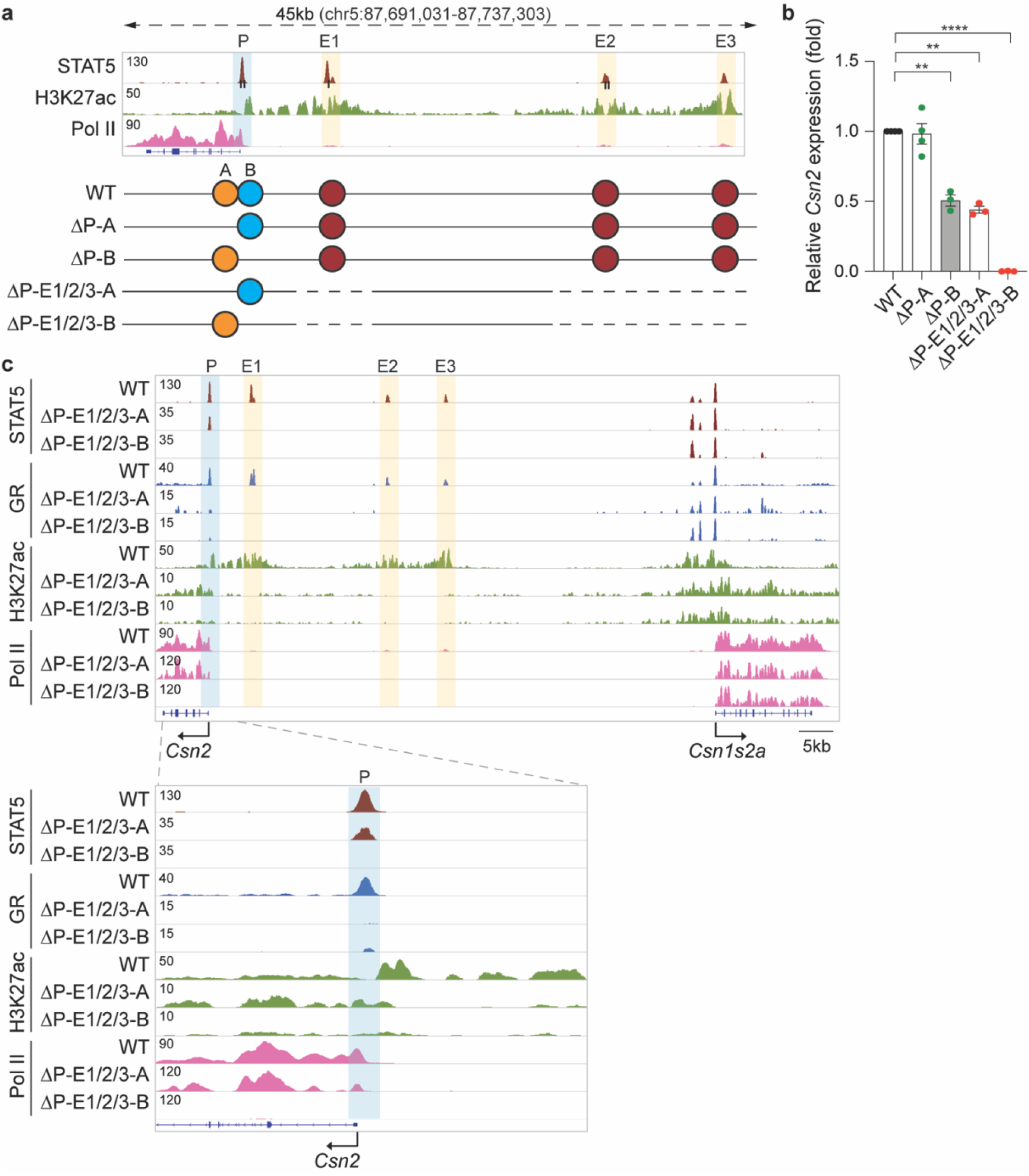
The non-canonical, but not the canonical, STAT5 site is required for *Csn2* promoter activity. **a,** Diagram of the *Csn2* promoter mutations introduced into the genome of WT and ΔE1/2/3 mice using base editing. The canonical GAS motif A is shown as orange circles, and the non-canonical GAS motif B is shown in blue. **b,** *Csn2* mRNA levels in lactating mammary tissues from WT and mutant mice were measured by qRT–PCR and normalized to *Gapdh* levels. Results are shown as the means ± s.e.m. of independent biological replicates (WT, ΔP-A, *n* = 4; ΔP-B, ΔP-E1/2/3-A, ΔP-E1/2/3-B, *n* = 3). One-way ANOVA followed by Dunnett’s multiple comparisons test was used to evaluate the statistical significance of differences between WT and each mutant mouse line. **c,** The *Csn2* locus including *Csn2* promoter was profiled using ChIP-seq in WT and mutant tissue.

**Fig. 5.**
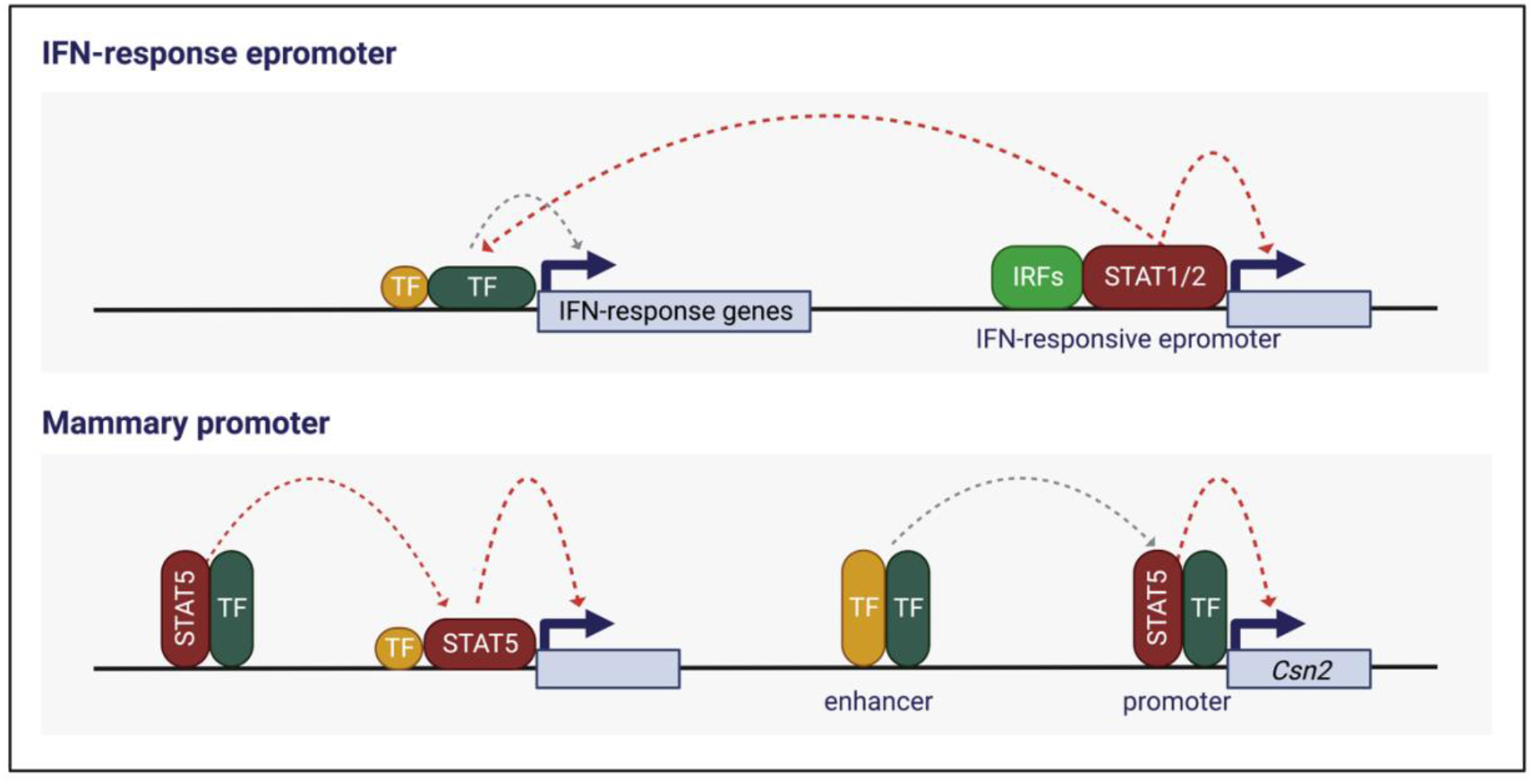
Model of the mammary Csn2 gene regulation. The *Csn2* gene is characterized by distinct enhancer and promoter elements, both harboring STAT5 binding sites (GAS motifs) that are bound by STAT5 during lactation. Individually, neither the enhancers nor the promoter STAT5 sites are required for efficient gene activation suggesting that they can compensate for each other’s activity. Inactivation of the distal enhancers and the promoter STAT5 site completely abrogates Csn2 expression suggesting that these regulatory elements function synergistically during pregnancy and lactation. Also, neighboring mammary genes are not impacted by the *Csn2* enhancer and promoter elements. Epromoters (reference) have been identified at interferon-response genes (reference) and they also harbor STAT binding sites but their regulation is distinctly different. In addition to their own native gene, Epromoters also control expression of neighboring genes. Neither Epromoter associated genes nor neighboring genes under their control harbor classical distal enhancers.

## Discussion

Based on our results of six genes^2^,^4^ (Lee et al., accompanying manuscript) uniquely expressed at very high levels in mammary tissue, a picture emerges that no unifying strategy can account for the extraordinary activation of genes during pregnancy and lactation, not even within the casein gene family that arose by gene duplication. While *Wap* gene expression is controlled by a single three-partite SE^2^, the five casein genes are subject to at least two different mechanisms. *Csn3* and *Csn1s2b* expression is largely controlled by a shared SE and gene-specific distal enhancers (Lee et al., accompanying manuscript). In contrast, regulation of the *Csn2* gene is unique in that it is under minimal influence of the SE and the distal enhancers by themselves are largely dispensable. Pregnancy-induced expression rather relies on the combined presence with a single cytokine-response promoter element and three distal enhancers. The *Csn2* gene is a unique and distinct example of distal enhancers and the promoter being controlled by the same set of cytokine response elements that bind the TF STAT5 and other mammary TFs.

Recent studies have identified a new class of regulatory elements, named Epromoters^9,10^, that function as both promoter and enhancer. Epromoters are hubs for TF machinery and typically associated with stress-response genes, such as those activated by interferons (IFN) through the TFs STAT1/2. A key feature of Epromoters is their capacity to activate neighboring genes independent of the presence of additional enhancers^10^. Epromoters are frequently found in loci harboring co-regulated genes, such as the *Oas* locus, that otherwise do not contain enhancers^10^.

Despite some similarities between Epromoters and the *Csn2* promoter element identified in this study, there are distinctive differences. Although both Epromoters and the five *casein* gene promoters, including *Csn2* investigated in this study, harbor cytokine-response elements bound by STAT TFs, their positioning appears to be different, with those in the casein genes being within 100 bp of the TSS and therefore likely integral part of the promoter. A distinct difference between the two regulatory elements is the capacity of Epromoters to activate neighboring genes in the absence of additional enhancers while the *Csn2* promoter element is fully dependent on the presence of distal enhancers. Moreover, unlike Epromoters, which activate neighboring genes at great distance, the *Csn2* promoter activity is confined to its own gene and neighboring genes are regulated independently. The activity of Epromoters has been validated in cell lines using CRISPR-Cas9 induced deletions spanning several hundred base pairs^10^, which leaves the possibility that additional elements contributed to their activity. In contrast, our study introduced single base pair mutations into GAS motifs demonstrating the specific and defining requirement of STAT5 in the activation of *Csn2.* Validating specificity of TF binding through the introduction of point mutations is desirable since it has been shown that phantom TF binding occurs at active promoters^11^.

Gene families regulated by IFNs, and milk protein genes share features that likely co-evolved during evolution. Repurposing structural genes, promoters and enhancers are a driving force in evolution, fostering innovation and the establishment of new genes and regulatory concepts^12,13^. Both, IFN-regulated genes and milk protein genes, are rapidly induced by cytokines that specifically utilize the JAK/STAT regulatory machinery permitting a rapid induction of genes. However, while transcriptional activation elicited by IFNs largely relies on JAK1 and STAT1/2^14^, mammary genes are predominantly activated by the pregnancy hormone prolactin, signaling through JAK2 and STAT5^6–8^. Gene expansion has significantly contributed to the evolution of both the IFN system and mammary genes, specifically the locus that harbors the five casein genes. Unlike the expansion that yielded possibly hundreds of IFN-responsive genes, gene duplication leading to milk production was limited, yet faced different challenges. Successful lactation demanded the acquisition of high-capacity regulatory building blocks, such as super-enhancers, that activated gene expression several thousand-fold during pregnancy, permitting five genes to account for 80% of milk protein.

Overall, our study suggests that the acquisition of promoter and enhancer elements, both responding to prolactin through the JAK/STAT pathway, was critical for the extraordinary expression levels of milk protein genes needed during lactation. The high-density of regulatory elements within the 330 kbp casein locus, including at least 15 enhancers and seven gene promoters, is a testimony to nature’s approach to build exceptionally active loci. Finally, our findings have significant implications for the mechanistic understanding of the evolution and function of regulatory elements rapidly responding to different cytokines yet depending on the same class of transcription factors.

## Materials and Methods

### Mice

All animals were housed and handled according to the Guide for the Care and Use of Laboratory Animals (8th edition) and all animal experiments were approved by the Animal Care and Use Committee (ACUC) of National Institute of Diabetes and Digestive and Kidney Diseases (NIDDK, MD) and performed under the NIDDK animal protocol K089-LGP-20. CRISPR/Cas9 targeted mice were generated using C57BL/6N mice (Charles River) by the transgenic core of the National Heart, Lung, and Blood Institute (NHLBI). Single-guide RNAs (sgRNA) were obtained from either OriGene (Rockville, MD) or Thermo Fisher Scientific (Supplementary Table 4). Target-specific sgRNAs and *in vitro* transcribed *Cas9* and *Base editor* mRNA were co-microinjected into the cytoplasm of fertilized eggs for founder mouse production. The ΔE1/2/3 and ΔP-E1/2/3 mutant mouse was generated by injecting a sgRNA for E1 into zygotes collected from ΔE2/3 mutant mice and for P into zygotes collected from ΔE1/2/3 mutant mice, respectively. All mice were genotyped by PCR amplification and Sanger sequencing (Quintara Biosciences) with genomic DNA from mouse tails (Supplementary Table 5).

### Chromatin immunoprecipitation sequencing (ChIP-seq) and data analysis

Mammary tissues from specific stages during pregnancy and lactation were harvested, and stored at −80°C. The frozen-stored tissues were ground into powder in liquid nitrogen. Chromatin was fixed with formaldehyde (1% final concentration) for 15 min at room temperature, and then quenched with glycine (0.125 M final concentration). Samples were processed as previously described^15^. The following antibodies were used for ChIP-seq: STAT5A (Santa Cruz Biotechnology, sc-1081 and sc-271542), GR (Thermo Fisher Scientific, PA1-511A), MED1 (Bethyl Laboratory, A300-793A), H3K27ac (Abcam, ab4729), RNA polymerase II (Abcam, ab5408), H3K4me1 (Active Motif, 39297) and H3K4me3 (Millipore, 07-473). Libraries for next-generation sequencing were prepared and sequenced with a HiSeq 2500 or 3000 instrument (Illumina). Quality filtering and alignment of the raw reads was done using Trimmomatic^16^ (version 0.36) and Bowtie^17^ (version 1.1.2), with the parameter ‘-m 1’ to keep only uniquely mapped reads, using the reference genome mm10. Picard tools (Broad Institute. Picard, http://broadinstitute.github.io/picard/. 2016) was used to remove duplicates and subsequently, Homer^18^ (version 4.8.2) and deepTools^19^ (version 3.1.3) software was applied to generate bedGraph files, seperately. Integrative Genomics Viewer^20^ (version 2.3.81) was used for visualization. Coverage plots were generated using Homer^18^ software with the bedGraph from deepTools as input. R and the packages dplyr (https://CRAN.R-project.org/package=dplyr) and ggplot2^21^ were used for visualization. Each ChIP-seq experiment was conducted for two replicates.

### Total RNA sequencing (Total RNA-seq) and data analysis

Total RNA was extracted from frozen mammary tissue from wild-type mice at day six of pregnancy and purified with RNeasy Plus Mini Kit (Qiagen, 74134). Ribosomal RNA was removed from 1 μg of total RNAs and cDNA was synthesized using SuperScript III (Invitrogen). Libraries for sequencing were prepared according to the manufacturer’s instructions with TruSeq Stranded Total RNA Library Prep Kit with Ribo-Zero Gold (Illumina, RS-122-2301) and paired-end sequencing was done with a HiSeq 3000 instrument (Illumina).

Total RNA-seq read quality control was done using Trimmomatic^16^ (version 0.36) and STAR RNA-seq^22^ (version STAR 2.5.3a) using paired-end mode was used to align the reads (mm10). HTSeq^23^ was to retrieve the raw counts and subsequently, R (https://www.R-project.org/), Bioconductor^24^ and DESeq2^21^ were used. Additionally, the RUVSeq^25^ package was applied to remove confounding factors. The data were pre-filtered keeping only those genes, which have at least ten reads in total. Genes were categorized as significantly differentially expressed with an adjusted p-value below 0.05 and a fold change > 2 for up-regulated genes and a fold change of < −2 for down-regulated ones. The visualization was done using dplyr (https://CRAN.R-project.org/package=dplyr) and ggplot2^26^.

### RNA isolation and quantitative real-time PCR (qRT–PCR)

Total RNA was extracted from frozen mammary tissue of wild type and mutant mice using a homogenizer and the PureLink RNA Mini kit according to the manufacturer’s instructions (Thermo Fisher Scientific). Total RNA (1 μg) was reverse transcribed for 50 min at 50°C using 50 μM oligo dT and 2 μl of SuperScript III (Thermo Fisher Scientific) in a 20 μl reaction. Quantitative real-time PCR (qRT-PCR) was performed using TaqMan probes (*Csn2*, Mm04207885_m1; *Csn1s2a,* Mm00839343_m1; mouse *Gapdh,* Mm99999915_g1, Thermo Fisher scientific) on the CFX384 Real-Time PCR Detection System (Bio-Rad) according to the manufacturer’s instructions. PCR conditions were 95°C for 30s, 95°C for 15s, and 60°C for 30s for 40 cycles. All reactions were done in triplicate and normalized to the housekeeping gene *Gapdh.* Relative differences in PCR results were calculated using the comparative cycle threshold (*C_τ_*) method and normalized to *Gapdh* levels. Results are shown as the means ± s.e.m. of independent biological replicates. A *t*-test or ANOVA were used to evaluate the statistical significance of differences between WT and mutant mice.

### Statistical analyses

All samples that were used for qRT–PCR, ChIP-seq, RNA-seq, 4C-seq and 3C were randomly selected, and blinding was not applied. For comparison of samples, data were presented as standard deviation in each group and were evaluated with a *t-*test and 2-way ANOVA multiple comparisons using PRISM GraphPad. Statistical significance was obtained by comparing the measures from wild-type or control group, and each mutant group. A value of \**P* < 0.05, \*\**P* < 0.001, \*\*\**P* < 0.0001, \*\*\*\**P* < 0.00001 was considered statistically significant. ns, no significant.

## Data availability

All data were obtained or uploaded to Gene Expression Omnibus (GEO). ChIP-seq and RNA-seq data from mutant mice in the study will be uploaded in GEO before publishing the manuscript. ChIP-seq and RNA-seq data of wild-type tissue at L1 and L10 were obtained under GSE74826, GSE119657, GSE115370 and GSE127144. Reviewer link will be shared upon request.

## Acknowledgments

We thank Ilhan Akan, Sijung Yun and Harold Smith from the NIDDK genomics core for NGS. This work utilized the computational resources of the NIH HPC Biowulf cluster (http://hpc.nih.gov). This work was supported by the Intramural Research Programs (IRPs) of National Institute of Diabetes and Digestive and Kidney Diseases (NIDDK) and National Heart, Lung, and Blood Institute (NHLBI).

## Author contributions

H. K.L. and L.H. designed the study. C.L. generated mutant mice. H.K.L. established mutant mouse line. H.K.L. performed experiments and data analysis. H.K.L. and L.H. supervised the study. H.K.L. and L.H. wrote the manuscript and all authors approved the final version.

## References

1. Panigrahi, A. & O’Malley, B.W. Mechanisms of enhancer action: the known and the unknown. Genome Biol 22, 108 (2021).

2. Shin, H.Y. et al. Hierarchy within the mammary STAT5-driven Wap super-enhancer. Nat Genet 48, 904–911 (2016).

3. Lee, H.K., Willi, M., Shin, H.Y., Liu, C. & Hennighausen, L. Progressing super-enhancer landscape during mammary differentiation controls tissue-specific gene regulation. Nucleic Acids Res 46, 10796–10809 (2018).

4. Lee, H.K., Willi, M., Kuhns, T., Liu, C. & Hennighausen, L. Redundant and non-redundant cytokine-activated enhancers control Csn1s2b expression in the lactating mouse mammary gland. Nat Commun 12, 2239 (2021).

5. Yamaji, D., Kang, K., Robinson, G.W. & Hennighausen, L. Sequential activation of genetic programs in mouse mammary epithelium during pregnancy depends on STAT5A/B concentration. Nucleic Acids Res 41, 1622–36 (2013).

6. Cui, Y. et al. Inactivation of Stat5 in mouse mammary epithelium during pregnancy reveals distinct functions in cell proliferation, survival, and differentiation. Mol Cell Biol 24, 8037–47 (2004).

7. Shillingford, J.M. et al. Jak2 is an essential tyrosine kinase involved in pregnancy-mediated development of mammary secretory epithelium. Mol Endocrinol 16, 563–70 (2002).

8. Liu, X. et al. Stat5a is mandatory for adult mammary gland development and lactogenesis. Genes Dev 11, 179–86 (1997).

9. Dao, L.T.M. et al. Genome-wide characterization of mammalian promoters with distal enhancer functions. Nat Genet 49, 1073–1081 (2017).

10. Santiago-Algarra, D. et al. Epromoters function as a hub to recruit key transcription factors required for the inflammatory response. Nat Commun 12, 6660 (2021).

11. Jain, D., Baldi, S., Zabel, A., Straub, T. & Becker, P.B. Active promoters give rise to false positive ‘Phantom Peaks’ in ChIP-seq experiments. Nucleic Acids Res 43, 6959–68 (2015).

12. Carelli, F.N., Liechti, A., Halbert, J., Warnefors, M. & Kaessmann, H. Repurposing of promoters and enhancers during mammalian evolution. Nat Commun 9, 4066 (2018).

13. Majic, P. & Payne, J.L. Enhancers Facilitate the Birth of De Novo Genes and Gene Integration into Regulatory Networks. Mol Biol Evol 37, 1165–1178 (2020).

14. Borden, E.C. et al. Interferons at age 50: past, current and future impact on biomedicine. Nat Rev Drug Discov 6, 975–90 (2007).

15. Metser, G. et al. An autoregulatory enhancer controls mammary-specific STAT5 functions. Nucleic Acids Res 44, 1052–63 (2016).

16. Bolger, A.M., Lohse, M. & Usadel, B. Trimmomatic: a flexible trimmer for Illumina sequence data. Bioinformatics 30, 2114–20 (2014).

17. Langmead, B., Trapnell, C., Pop, M. & Salzberg, S.L. Ultrafast and memory-efficient alignment of short DNA sequences to the human genome. Genome Biol 10, R25 (2009).

18. Heinz, S. et al. Simple combinations of lineage-determining transcription factors prime cis-regulatory elements required for macrophage and B cell identities. Mol Cell 38, 576–89 (2010).

19. Ramirez, F. et al. deepTools2: a next generation web server for deep-sequencing data analysis. Nucleic Acids Res 44, W160–5 (2016).

20. Thorvaldsdottir, H., Robinson, J.T. & Mesirov, J.P. Integrative Genomics Viewer (IGV): high-performance genomics data visualization and exploration. Brief Bioinform 14, 178–92 (2013).

21. Love, M.I., Huber, W. & Anders, S. Moderated estimation of fold change and dispersion for RNA-seq data with DESeq2. Genome Biol 15, 550 (2014).

22. Dobin, A. et al. STAR: ultrafast universal RNA-seq aligner. Bioinformatics 29, 15–21 (2013).

23. Anders, S., Pyl, P.T. & Huber, W. HTSeq--a Python framework to work with high-throughput sequencing data. Bioinformatics 31, 166–9 (2015).

24. Huber, W. et al. Orchestrating high-throughput genomic analysis with Bioconductor. Nat Methods 12, 115–21 (2015).

25. Risso, D., Ngai, J., Speed, T.P. & Dudoit, S. Normalization of RNA-seq data using factor analysis of control genes or samples. Nat Biotechnol 32, 896–902 (2014).

26. Wickham, H. Ggplot2: elegant graphics for data analysis, viii, 212 p. (Springer, New York, 2009).

